# Fast, volumetric live-cell imaging using high-resolution light-field microscopy

**DOI:** 10.1101/439315

**Authors:** Haoyu Li, Changliang Guo, Deborah Kim-Holzapfel, Weiyi Li, Yelena Altshuller, Bryce Schroeder, Wenhao Liu, Yizhi Meng, Jarrod B. French, Ken-Ichi Takamaru, Michael A. Frohman, Shu Jia

## Abstract

Visualizing diverse anatomical and functional traits that span many spatial scales with high spatio-temporal resolution provides insights into the fundamentals of living organisms. Light-field microscopy (LFM) has recently emerged as a scanning-free, scalable method that allows for high-speed, volumetric functional brain imaging. Given those promising applications at the tissue level, at its other extreme, this highly-scalable approach holds great potential for observing structures and dynamics in single-cell specimens. However, the challenge remains for current LFM to achieve subcellular level, near-diffraction-limited 3D spatial resolution. Here, we report high-resolution LFM (HR-LFM) for live-cell imaging with a resolution of 300-700 nm in all three dimensions, an imaging depth of several micrometers, and a volume acquisition time of milliseconds. We demonstrate the technique by imaging various cellular dynamics and structures and tracking single particles. The method may advance LFM as a particularly useful tool for understanding biological systems at multiple spatio-temporal levels.

## 1. Introduction

Light-field microscopy (LFM) simultaneously captures both the 2D spatial and 2D angular information of the incident light, allowing computational reconstruction of the full 3D volume of a specimen from a single camera frame [1–4]. Conventionally, fluorescent imaging techniques acquire 3D spatial information in a sequential or scanning fashion [5–12], inevitably compromising temporal resolution and increasing photodamage for live imaging. The 4D imaging scheme of LFM effectively liberates volume acquisition time (limited primarily by the camera’s frame rate) from the spatial parameters (e.g. the field of view (FOV) and spatial resolution), thus making LFM a promising tool for high-speed, volumetric imaging of living biological systems with low photodamage across many spatial levels.

Towards the tissue level, the latest LFM techniques have demonstrated promising results for functional brain imaging with cellular level spatial resolution of several micrometers across a depth of tens to hundreds of micrometers, and at a fast volume acquisition time on the order of 10 milliseconds [4,13–15]. At its other extreme, this highly-scalable approach holds great potential for visualizing delicate structures and dynamics within single-cell specimens. However, the challenge remains for current LFM to achieve subcellular level, near-diffraction-limited 3D spatial resolution.

For the existing LFM techniques, in practice, a microlens array (MLA) is placed at the native image plane (NIP) of a wide-field microscope, and the optical signal is recorded in an aliased manner across the microlenses at the back focal plane of the MLA [1,2]. The development of wave-optics models allows high-resolution reconstruction of densely aliased high-spatial frequencies through point-spread function (PSF) deconvolution [3,4]. However, near the NIP, the sampling pattern of the spatial information becomes redundant, resulting in prohibitive artifacts [3,4]. This limitation causes non-uniform resolution across depth, hindering applications that require visualization of fine 3D information spreading over a large axial range in a sample. Current LFM techniques circumvent the artifacts mainly by imaging on one side of the NIP [3,4,14] or using a coded wavefront [16]. These approaches are feasible for functional brain imaging (e.g. using 20× or 40× objective lenses) despite compromised depth of focus (DOF) or spatial resolution. However, they may be prohibitive for high-resolution microscopy, because they would either sacrifice the already tightened DOF (typically <1 μm for a 100×, 1.45NA oil objective lens) or worsen the 3D spatial resolution away from the diffraction limit. Alternatively, the MLA may instead form an imaging relationship between the NIP and the camera sensor, as used in the focused plenoptic camera [17]. While the spatial information at the NIP can thereby be recovered using this scheme, the sampling geometry of the angular information becomes less aliased and redundant, inherently impairing the refocusing (or volumetric imaging) capability for practical applications. Hence, new optical design and computational framework are highly desired for high-resolution light-field imaging in single-cell specimens.

Here, we develop a HR-LFM method for live-cell imaging by simultaneous, dense sampling of both spatial and angular information. HR-LFM achieves a 3D spatial resolution of 300-700 nm, an imaging depth of several micrometers, and a volume acquisition time down to milliseconds. We demonstrated the technique by imaging various cellular systems and tracking single particles.

## Methods

### 2.1 Experimental Setup

We constructed HR-LFM on an epifluorescence microscope (Nikon Eclipse Ti-U) using a 100×, NA=1.45, oil immersion objective lens (Nikon CFI-PLAN 100×, 1.45 NA) (**Fig. 1a** and **Appendix 1**). The sample stage was controlled by a nano-positioning system (Prior). The samples were illuminated with 488-nm, 561-nm and 647-nm lasers (MPB Communications). The corresponding emitted fluorescence was collected using dichroic mirrors (T495lpxr, T560lpxr and T660lpxr, Chroma, respectively) and emission filters (ET525/50, Chroma; FF02-617/73, Semrock; ET700/75, Chroma, respectively). A NA-matched MLA (S125-F30, RPC Photonics) was aligned in a five-axis kinematic mount (K5X1, Thorlabs). The light field was imaged using a 1:1 relay lens (Nikon AF-S VR Micro-Nikkor 105mm f/2.8G IF-ED) and recorded on a scientific complementary metal-oxide-semiconductor (sCMOS) camera (ORCA-Flash4.0, Hamamatsu).

In the setup, the MLA forms a defocused imaging relationship as 1/*a* + 1/*b* > 1/*f*_*ml*_, where *a* and *b* denote the distances to the NIP and the camera sensor, respectively, and *f*_*ml*_ is the focal length of the MLA. This design contrasts with conventional LFM [1] (*a* = 0 and *b* = *f*_*ml*_) and the focused plenoptic camera [17] (1/*a* + 1/*b* = 1/*f*_*ml*_) (**Appendix 2**), and benefits HR-LFM in two main ways. First, the distance *a* can displace the artifact region away from the DOF by *a*/*M*^2^ in the object space, where *M* is the magnification of the objective lens. Second, the distance *b* facilitates optimum dense sampling of both the spatial and the angular information of optical signals on the camera sensor. In this work, the values of *a* and *b* (*a* = 25 mm and *b* = 4 mm) were carefully determined using numerical simulations to obtain the optimized spatio-angular distribution for reconstruction of point-emitters distributed in the 3D space (**Appendix 3**, **Eqs. 1-5**). Using this scheme, the optical signals imaged by the objective lens can be densely aliased compared to conventional LFM, capturing different perspectives of the sample onto the camera sensor (**Fig. 1b-d**). The aliasing of the recorded data can thus effectively suppress the reconstruction artifacts at the NIP, substantially improving the DOF and spatial resolution.

**Fig. 1.**
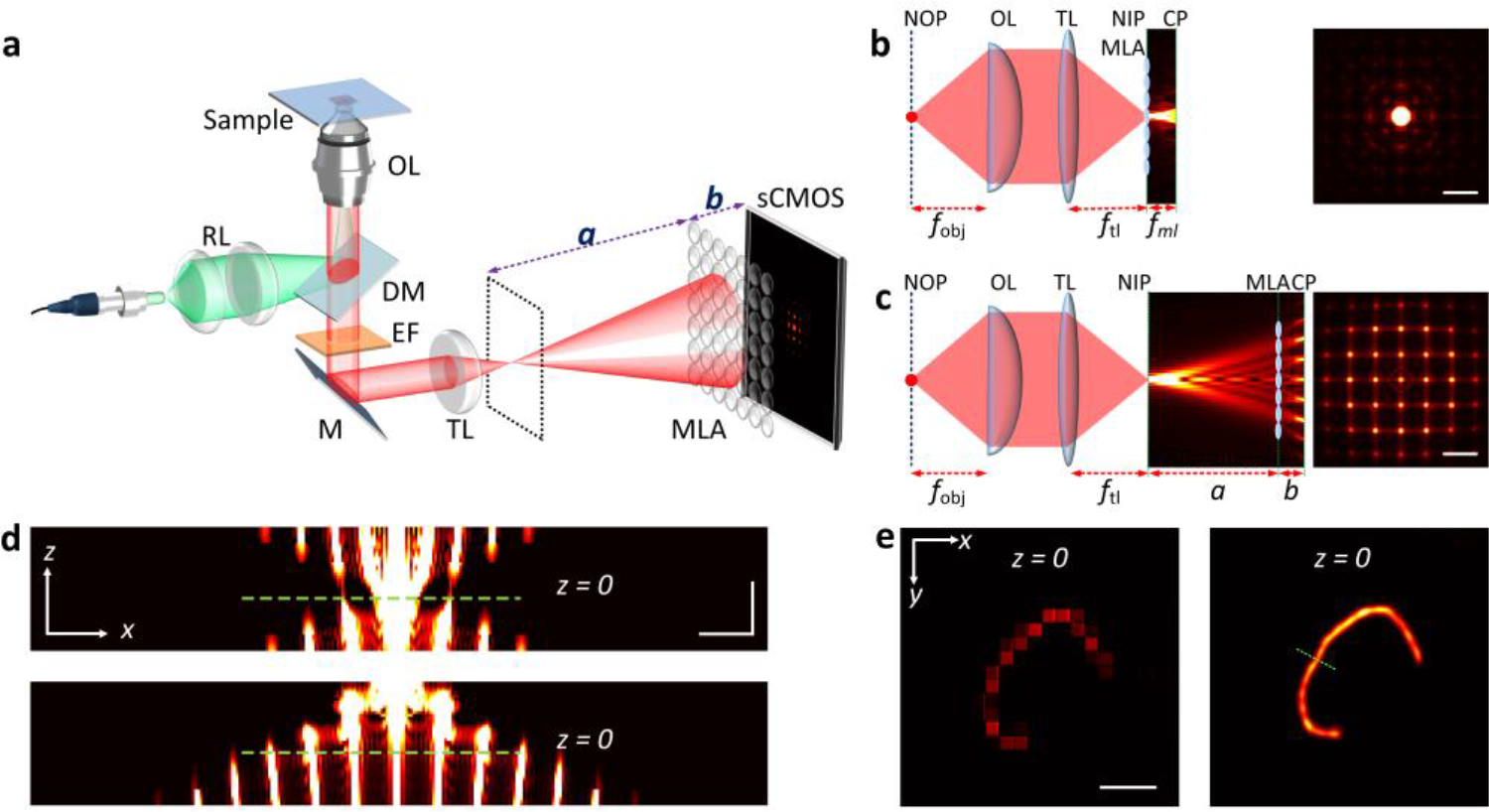
High-resolution light-field microscopy (HR-LFM). (a) A schematic of the experimental setup for HR-LFM. The objective lens (OL) and tube lens (TL) form an image at the native image plane (NIP, dashed plane). The microlens array (MLA) is situated at *a* and *b* to the NIP and the sCMOS camera, respectively. RL: relay lenses; DM: dichroic mirror; EF: emission filter; M: mirror. (b,c) Left panel, simulated light propagation from a point emitter located at the native object plane (NOP) in conventional LFM (b) and HR-LFM (c). Right panel, *x*-*y* views of the corresponding light-field information captured at the camera plane (CP). *f*_obj_, the effective focal length of the objective lens; *f*_tl_, the focal length of the tube lens; *f*_*ml*_, the focal length of the MLA. (d) *x*-*z* views of the propagation of the optical signals in conventional LFM (top) and HR-LFM (bottom) from *z* = −2.0 μm to 2.5 μm. The dashed lines represent the NIPs. (e) Reconstruction of aggregated fluorescent beads placed at the NOP, using conventional LFM (left) and HR-LFM (right). Reconstruction artifacts were observed in conventional LFM. The profile across the dashed line exhibits FWHM of 387 nm. Scale bars, 2 μm (b-d), 5 μm (e).

### 2.2 Reconstruction Algorithm

To reconstruct the volumetric data, the Fresnel propagation of light by the distances of *a* and *b*, i.e. a defocused PSF, was established using the scalar diffraction theory [18]. Specifically, the final intensity image *o*(x″) at the camera plane is described by *o*(*x*″) = ⎰|*h*(x″, p)|^2^*g*(p)*d*p (**Appendix 3**, **Eq. 5**), where 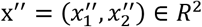 represents the real coordinates (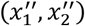) on the camera plane, p ∈ *R*^3^ are the real coordinates of a point source in a volume in the object domain, whose combined intensities are distributed according to *g*(p). *h*(x″, p) represents the complex-valued PSF, which considers, sequentially, the light propagation through the high-NA objective lens, Fresnel propagation of light by the distance of *a*, modulation induced by the MLA, and another Fresnel propagation to the camera plane by the distance of *b* (**Appendix 3**, **Eqs. 1-4**). In practice, considering the discrete model, *h*(x″, p) is represented by the measurement matrix *H*, which elements *h*_kj_ describe the projection of light arriving at the pixel *o*(*j*) on the camera plane from the *k*^th^ voxel *g*(*k*) in the object space. The volumetric information was then reconstructed employing the wave-optics model [3,4] based on an inverse-problem deconvolution framework [19] (**Appendix 3** and **Supplementary Code**). To verify the algorithm, we recorded light-field images of 100-nm fluorescent beads placed on the native object plane (NOP) of the objective lens, where the aggregated beads can be properly reconstructed on the NIP with near-diffraction-limited widths of ~400 nm without artifacts using HR-LFM, compared to conventional LFM (**Fig. 1e**).

### 2.3 System Characterization

**Fig. 2.**
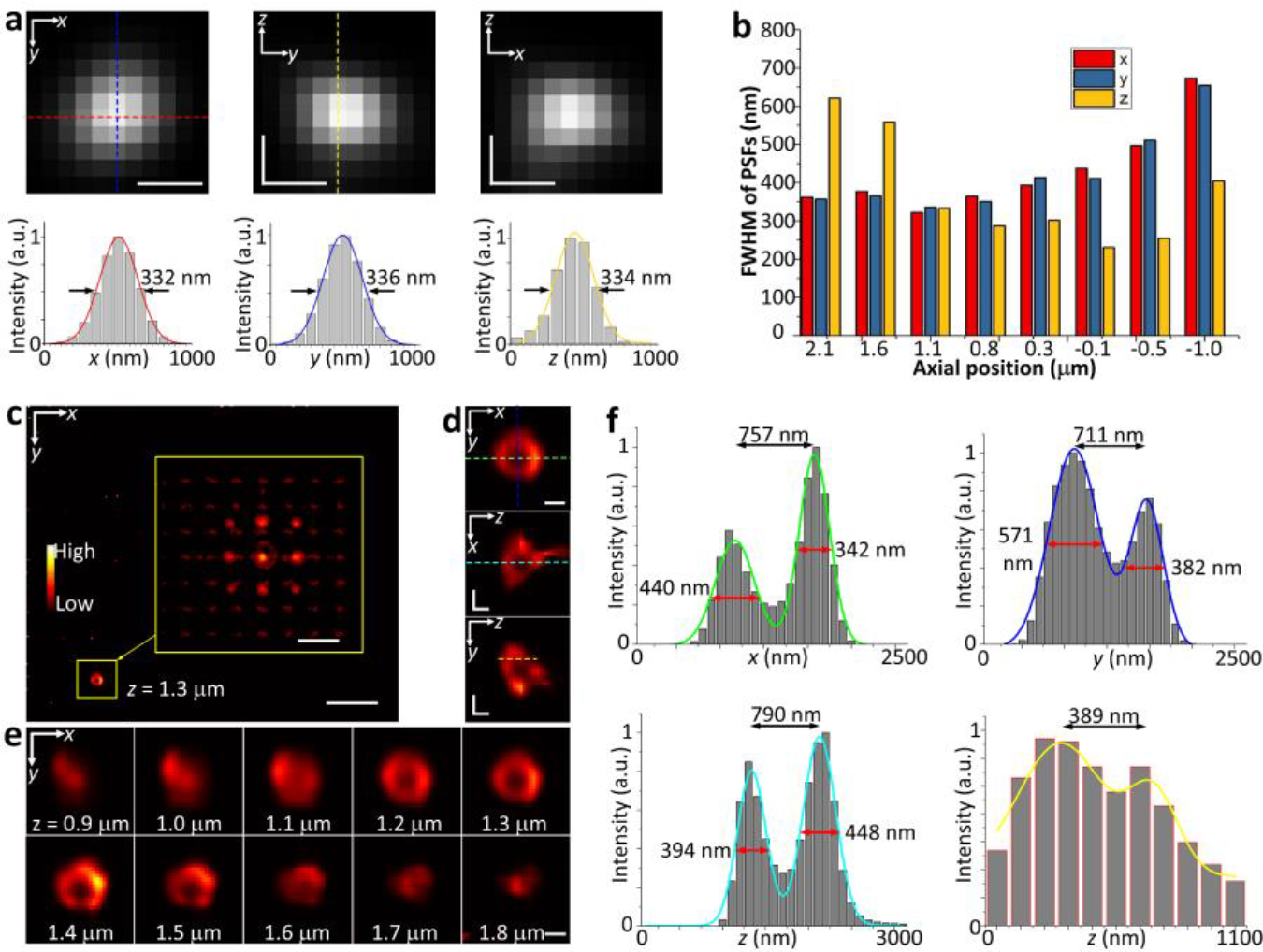
Characterization of HR-LFM. (a) The reconstructed cross-sectional images (top panel) and profiles (bottom panel) of 100-nm fluorescent beads in *x*-*y* (left), *y*-*z* (middle) and *x*-*z* (right) at *z* = 1.1 μm. The profiles along the dashed lines exhibit isotropic FWHMs of 332 nm, 336 nm, and 334 nm in *x*, *y*, and *z*, respectively. (b) The 3D FWHM values of the reconstructed PSFs at various axial positions over a >3-μm range. (c) A reconstructed *z*-stack image at *z* = 1.3 μm of a surface-stained, 1-μm fluorescent microsphere. The hollow structure was observed. Inset, raw HR-LFM data of the boxed microsphere taken at a volume acquisition time of 0.2 s. (d) Lateral (*x*-*y*) and axial (*x*-*z* and *y*-*z*) cross-sectional views of the reconstructed microsphere in (c). (e) The corresponding *z*-stack images of the microsphere at an axial step size of 100 nm. (f) Corresponding lateral and axial cross-sectional profiles along colored dashed lines in (d). The profiles exhibited the sub-micrometer hollow structure resolved by HR-LFM and the FWHMs of the stained surface of ~400 nm at *z* = 1.3 μm in all three dimensions. Scale bars, 300 nm (a), 5 μm (c), 2 μm ((c) inset), 500 nm (d,e).

Next, to characterize HR-LFM, we imaged 100 nm fluorescent beads and measured the reconstructed PSFs of the system at varying depths (**Fig. 2a,b**, **Appendices 4** and **5**). The numbers of iterations taken for the reconstruction at varying depths were determined based on the distribution of the optical signals across the MLA and the corresponding signal-to-noise ratio (SNR). These numbers were consistently used in this work for other samples (**Appendix 4**). The full-width at half-maximum (FWHM) values of these reconstructed PSFs at each depth exhibited a near-diffraction-limited 300-700 nm resolution in all three dimensions over a >3 μm range, ~4× larger than the corresponding DOF of wide-field microscopy. Detailed features of the system are shown in **Appendix 4**. First, when the beads were scanned near the axial position *z* = *a*/*M*^2^ = 2.5 μm (*a* = 25 mm and *M* = 100; i.e. forming images near the MLA), the optical signals were mainly restrained into a single microlens as in conventional LFM. Thus, HR-LFM performed as a NA-matched single-lens imaging system, exhibiting spatial resolution of ~300 nm and 600 nm in the lateral and axial dimensions, respectively, consistent with the PSF measured in wide-field microscopy using the same 100×, 1.45NA objective lens. Second, when the beads were scanned toward the focal plane *z* = 0 (i.e. forming images between the MLA and the NIP), the optical signals became broadly distributed on the MLA, which led to higher angular sensitivity, thereby improving the axial resolution (**Appendix 5**). Notably, the lateral resolution was steadily maintained using the wave-optics model, and thus the system achieved a near-isotropic 3D resolution within this axial range. Lastly, beyond the NIP (*z* < 0), the optical signals became further broadened on the MLA due to the diffraction of light. While the high axial sensitivity was maintained, the lateral resolution became worse due to the degraded signal detection from an increased number of microlenses. In general, the implementation of wave-optics based reconstruction effectively overcame the tradeoff and maintained a high resolution in both the axial and lateral dimensions. Furthermore, the use of the MLA permits sensitive angular detection of the wavefront (i.e. spatial frequencies), improving axial-resolving capability within a substantial range of the DOF. These features make HR-LFM distinct from a conventional diffraction-limited imaging system, providing a new type of optical engineering scheme for high-resolution microscopy.

## 3 Results

### 3.1 Imaging Caliber Structures and Fixed Biological Samples

Further measurements, in good agreement with the reconstructed PSF values, have been obtained using known caliber structures and fixed biological samples. We first imaged surface stained, 1-μm fluorescent microspheres (F14791, ThermoFisher), which sub-micrometer hollow structure was resolved using HR-LFM (**Fig. 2c-e**). The corresponding lateral and axial cross-sectional profiles exhibited FWHMs of the stained surface of 300-500 nm in all three dimensions (**Fig. 2f**), consistent with the measured values in **Fig. 2b** and **Appendix 4**.

Next, we imaged immuno-labeled mitochondria in HeLa cells. For sample preparation, HeLa cells were obtained from the American Type Culture Collection (ATCC), maintained in 1× Minimum Essential Medium (MEM) (Corning CellGro) with 10% fetal bovine serum (FBS) (Atlanta Biologicals) and 50 μg/ml gentamycin (Amresco), and incubated at 37°C with 5% CO2. The cells were plated on a 35 mm^2^ MatTek glass-bottom dishes (MatTek), incubated at 37°C for 16 hours, and fixed with 4% (vol/vol) formaldehyde (15735, Electron Microscopy Sciences) prepared in phosphate buffered saline (PBS) for 10 mins at 37°C. The cells were then blocked and permeabilized with blocking and antibody dilution buffer (1% (vol/vol) bovine serum albumin (BSA) (Santa Cruz Biotechnologies) and 0.25% (vol/vol) Triton X-100 prepared in PBS) for 1 hour at room temperature. The cells were then incubated with the mitochondrial marker primary antibody mouse anti-Tom20 (Santa Cruz Biotechnologies F10, SC-17764), at 1 μg/ml in blocking and antibody dilution buffer for 2 hours while gently shaking at room temperature. The cells were then washed 3 times with PBS for 5 mins each. Secondary antibody (AlexaFluor 647-conjugated AffiniPure Goat Anti-Mouse IgG, 1 mg/ml, Jackson ImmunoResearch) was diluted 1:1000 in 1% BSA in PBS and incubated with gentle shaking for 1 hour at room temperature. The cells were washed 3 times with PBS for 5 mins. The cells were placed in imaging buffer (20 mM HEPES pH 7.4, 135 mM NaCl, 5 mM KCl, 1 mM MgCl_2_, 1.8 mM CaCl_2_, 5.6 mM glucose) before imaging.

As seen, HR-LFM recorded the full 4D light-field information (**Fig. 3a**), allowing us to synthesize the focal stacks of the entire volume of the specimen (**Fig. 3b**). Remarkably, HR-LFM captured mitochondria that were out-of-focus and poorly detected by wide-field microscopy due to its limited DOF (**Fig. 3c** and **Appendix 6**). The mitochondrial structures were well-resolved by HR-LFM across a ~3-μm axial range in all three dimensions without the need for any sample or focal-plane scanning. Furthermore, as a comparison, conventional LFM can only image cells on one side of the NIP to avoid the prohibitive artifacts, resulting in substantially degraded volumetric imaging capability, especially in the axial dimension such as the DOF and axial resolution (**Fig. 3d-f**).

**Fig. 3.**
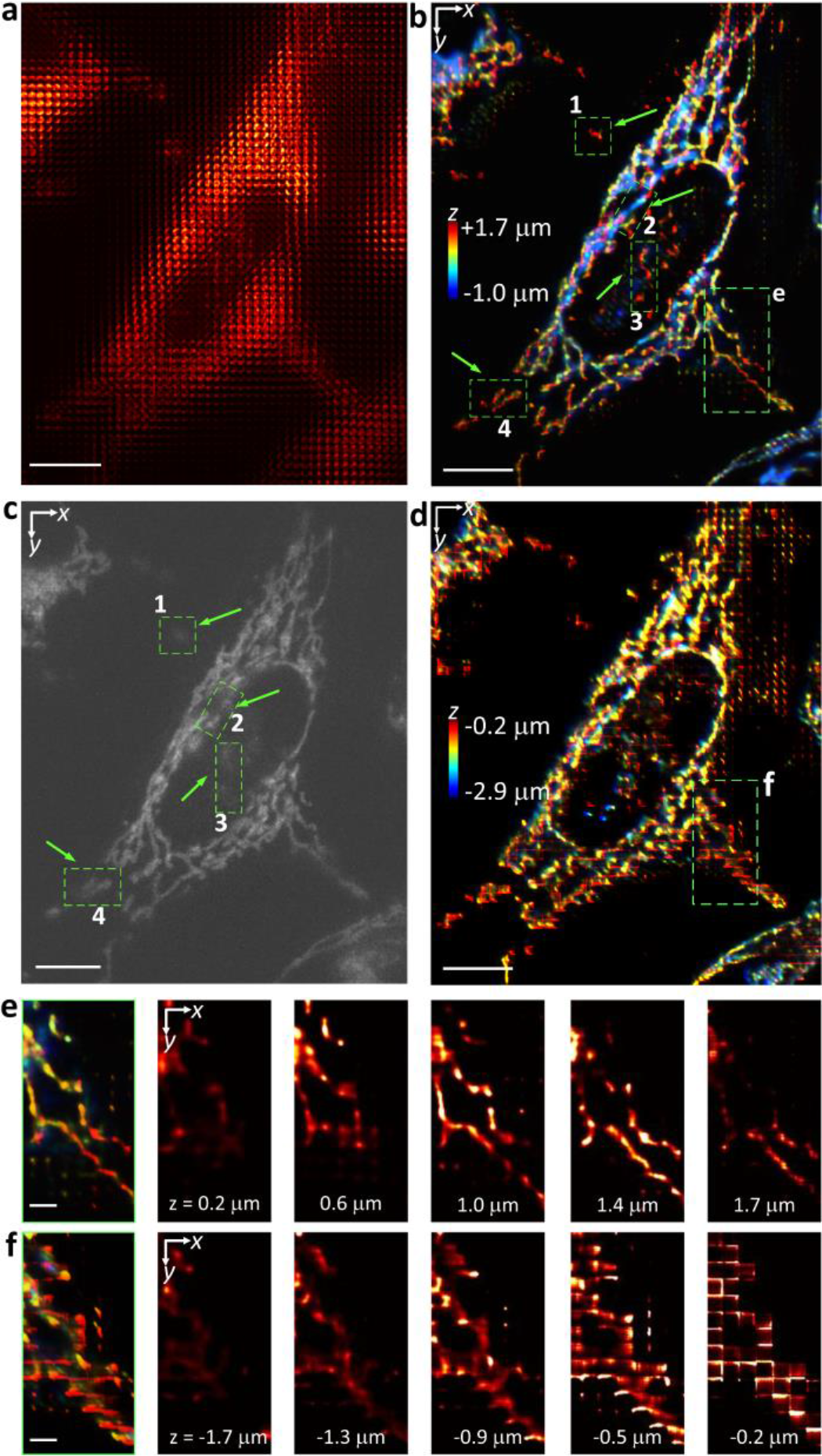
Imaging mitochondria in fixed HeLa cells using HR-LFM. (a-d) Raw light-field (a), reconstructed 3D HR-LFM (b), conventional wide-field (c) and conventional LFM (d) images of immuno-labeled mitochondria in HeLa cells. The depth-information is color-coded according to the color scale bars in (b,d). Boxed regions (1-4) in (b,c) show that HR-LFM captured mitochondria that were out-of-focus and hence poorly detected by conventional wide-field microscopy due to its limited DOF. (e,f) Zoomed-in images (leftmost panel) of the boxed regions in (b,d), respectively, and their corresponding selected *z*-stack images (right panel), showing both sensitive axial discrimination and suppression of reconstruction artifacts using HR-LFM. Scale bars, 10 μm (a-d), 2 μm (e,f).

Furthermore, we imaged the Golgi complex in HeLa cells, immuno-labeled for the Golgi marker GM130 with DyLight 549, where the nearby Golgi structures as close as ~400 nm in all three dimensions were resolved by HR-LFM (**Appendix 7**).

### 3.2 Imaging Mitochondria in Living Drp1^−/−^ Mouse Embryo Fibroblasts (MEFs)

To demonstrate live-cell imaging, we first recorded mitochondrial dynamics in living *Drp1*^−/−^ MEFs [20], labeled with MitoTracker, a mitochondrial fluorescent tracking dye (**Fig. 4**). The dynamin-related GTPase (*Drp1*) mediates mitochondrial division and distribution, playing a critical role in mammalian development [20]; cells lacking *Drp1* have low rates of mitochondrial fission and thus have long mitochondria.

**Fig. 4.**
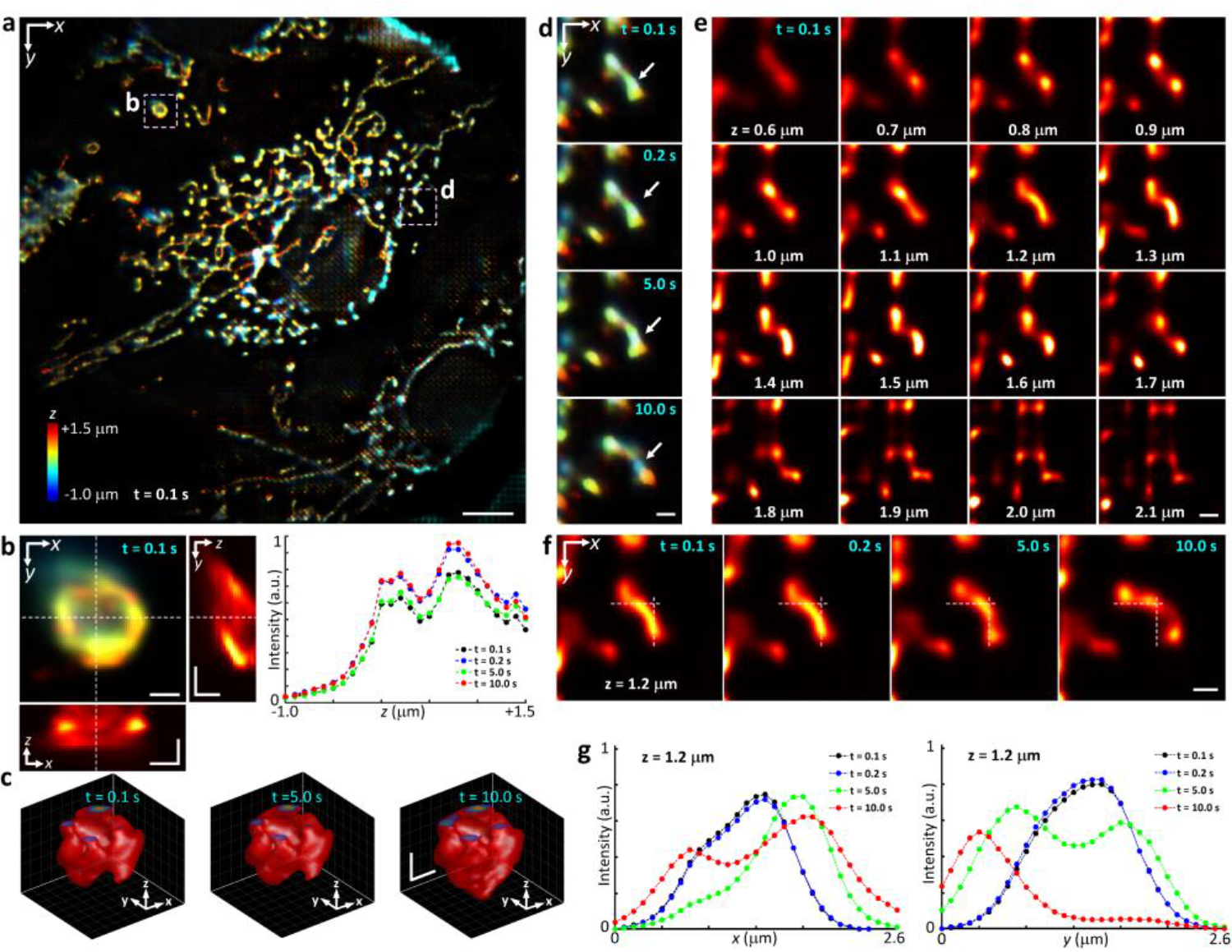
Imaging mitochondria in living *Drp1*^−/−^ mouse embryo fibroblasts (MEFs) using HR-LFM. (a) Reconstructed 3D image of mitochondria labeled with MitoTracker in living *Drp1*^−/−^ MEFs at time-point *t* = 0.1 s, taken at a volume acquisition time of 0.1 s. The depth-information is color-coded according to the color scale bar. (b) Left, zoomed-in image of the corresponding boxed region in (a) and cross-sectional *x*-*z* and *y*-*z* views along the dashed lines. Right, the corresponding axial cross-sectional profiles at time-points *t* = 0.1 s, 0.2 s, 5.0 s, and 10.0 s, showing resolved axial structures separated by ~500 nm. (c) Volume rendering of the same region in (b) at time-points *t* = 0.1 s, 5.0 s, and 10.0 s, respectively. (d) Zoomed-in images of the corresponding boxed region in (a) at time-points *t* = 0.1 s, 0.2 s, 5.0 s, and 10.0 s, respectively. White arrows indicate mitochondria undergoing structural reorganization. (e) The *z*-stack images of (d) from *z* = 0.6 μm to 2.1 μm at *t* = 0.1 s, showing axial structural changes captured at an axial step size of 100 nm. (f) The *z*-stack image of (d) at *z* = 1.2 μm at time-points *t* = 0.1 s, 0.2 s, 5.0 s, and 10.0 s, respectively. (g) The cross-sectional profiles along the dashed lines in (f) exhibit sub-micrometer structural changes resolved in *x* (left) and *y* (right) at time-points *t* = 0.1 s, 0.2 s, 5.0 s, and 10.0 s, respectively. Scale bars: 10 μm (a), 1 μm (b-f).

*Drp1*^−/−^ MEFs were generously provided by Hiromi Sesaki (Johns Hopkins University). For sample preparation, *Drp1*^−/−^ MEFs were cultured in DMEM + 10% FBS and plated into tissue culture dishes containing sterile coverslips and cultured for 1-2 days until 50-80% confluent. The cells were incubated in medium containing a 1:5000 dilution of MitoTracker Deep Red (M22426, ThermoFisher) for 30 mins at 37°C, washed 3 times with PBS, and then placed back into growth medium lacking MitoTracker.

Using the full sCMOS camera chip (2048 × 2048 pixels), we captured the entire FOV (>100 × 100 μm laterally and >3-5 μm axially) enclosing several cells at a volume acquisition time of 0.1 s (**Fig. 4a**). The high spatial resolution allows us to visualize fine mitochondrial structures and distributions as close as 500 nm in all three dimensions (**Fig. 4b-g**, **Visualization 1** and **Visualization 2**). Without the need for scanning, HR-LFM permits low light exposure (0.05-0.5 W cm^−2^) for time-lapse acquisition over more than thousands of time-points (up to minutes) without obvious photodamage or photobleaching [21]. We observed extension and retraction of mitochondrial tubules, where mitochondrial movements of up to 0.53 μm s^−1^ were recorded, as well as occasional mitochondrial division (**Visualization 1** and **Visualization 3**).

### 3.3 Imaging Golgi-derived Membrane Vesicles in Living COS-7 Cells

**Fig. 5.**
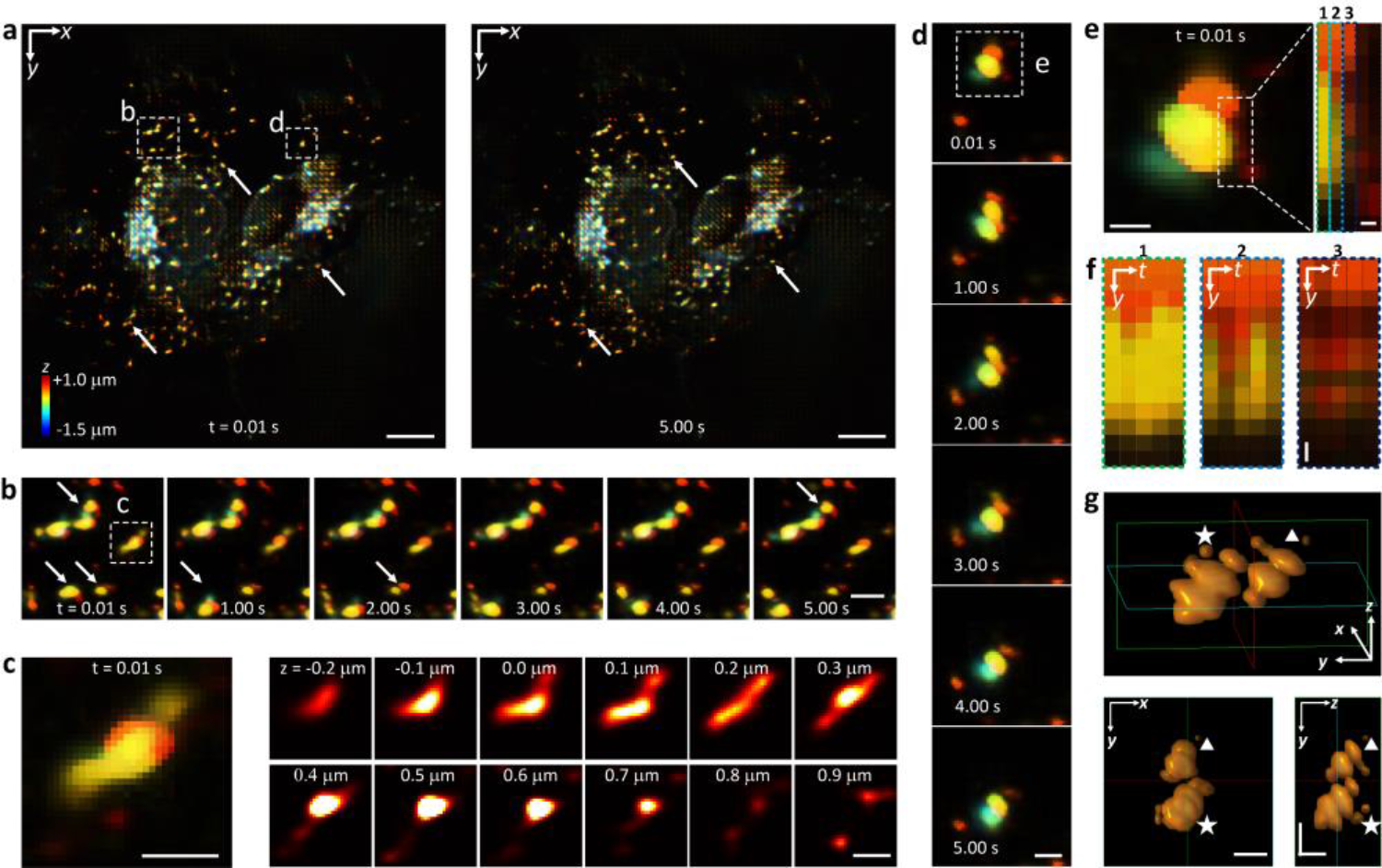
Imaging Golgi-derived membrane vesicles in living COS-7 cells using HR-LFM. (a) Reconstructed 3D images of Golgi-derived membrane vesicles labeled with mEmerald-Golgi-7 in living COS-7 cells at time-points *t* = 0.01 s (left) and 5.00 s (right), taken at a volume acquisition time of 0.01 s. The depth-information is color-coded according to the color scale bar. White arrows indicate multiple moving vesicles. (b) Zoomed-in images of the corresponding boxed region in (a) at time-points *t* = 0.01 s, 1.00 s, 2.00 s, 3.00 s, 4.00 s, and 5.00 s. White arrows indicate multiple moving vesicles. (c) Zoomed-in image (left panel) of the boxed region in (b) and its *z*-stack images (right panel) from *z* = −0.2 μm to 0.9 μm at an axial step size of 100 nm, showing axially-resolved structural variations of nearby vesicles. (d) Zoomed-in images of the corresponding boxed region in (a) at time-points *t* = 0.01 s, 1.00 s, 2.00 s, 3.00 s, 4.00 s, and 5.00 s. (e) Zoomed-in image of the boxed region in (d). The inset magnifies the boxed region in (e) spanning several vesicles. (f) Each *y*-*t* image (1-3) respectively shows the intensity variations of individual columns of pixels (1-3) in the inset of (e) over five consecutive time-points *t* = 0.01 s, 0.02 s, 0.03 s, 0.04 s, and 0.05 s. (g) 3D (top), *x*-*y* (bottom left) and *y*-*z* (bottom right) views of the moving vesicles in (e) at time-points *t* = 0.01 s (triangle) and 4.00 s (star), resolving vesicles separated as close as 300-500 nm in all three dimensions. Scale bars: 10 μm (a), 2 μm (b), 1 μm (c, d, g), 500 nm (e), 100 nm ((e) inset, f).

We next imaged Golgi-derived membrane vesicles in living COS-7 cells, labeled with a trans-Golgi marker, mEmerald-Golgi-7 (**Fig. 5** and **Appendix 8**). These vesicles occasionally undergo rapid movement between the Golgi complex and other intracellular organelles, posing a challenge for capturing their volumetric dynamics using scanning-based imaging methods.

For sample preparation, COS-7 cells were obtained from ATCC and grown in Dulbecco’s modified Eagle’s medium (DMEM) supplemented with 10% FBS and 100 U/ml penicillin streptomycin. Cells were seeded in a 35 mm^2^ MatTek glass-bottom dishes (MatTek), transfected with the trans-Golgi marker mEmerald-Golgi-7 (Addgene 54108), which carries the 1-82 aa residues of β-1,4-galactosyltransferase-I (β-1,4-GalT-I) at the N-terminus, using Lipofectamine 2000 (ThermoFisher), and imaged 24 hours post-transfection.

Using the full camera chip, we captured their rapid 3D motions at a volume acquisition time of 0.01 s over hundreds to thousands of camera frames (**Fig. 5a**). Notably, the high spatio temporal resolution allowed us to observe the rapid interactions of individual vesicles separated as close as 300-500 nm and moving up to 2.06 μm s^−1^ in all three dimensions (**Fig. 5b-g**and **Visualization 4**). In addition, the volume acquisition speed can be further accelerated by imaging only a region of interest on the camera chip. We imaged the same samples at a volume acquisition time faster than 5 ms without noticeable degradation in image quality or resolution (**Appendix 9**).

### 3.3 Tracking Hydrophobically-modified Glycol Chitosan (HGC) Nanoparticles

**Fig. 6.**
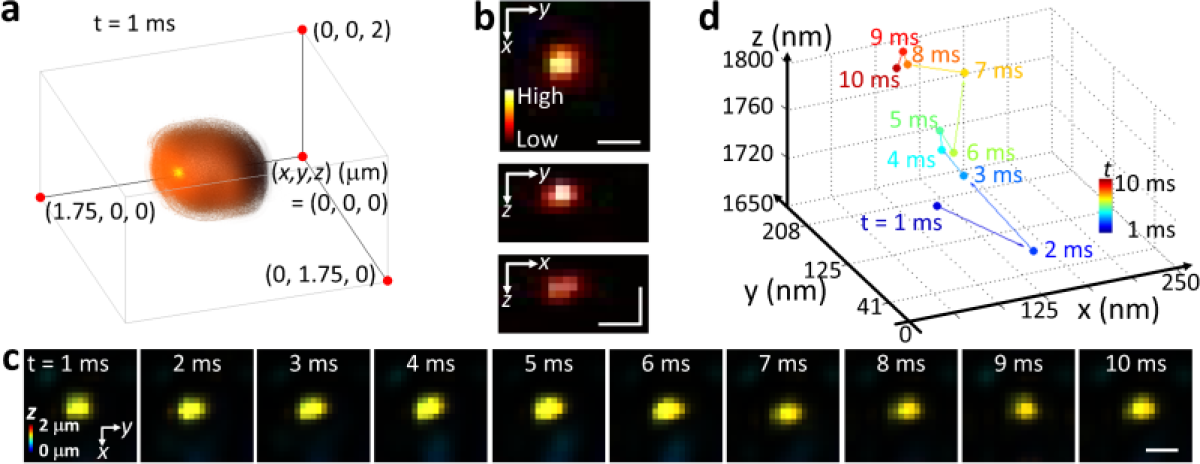
Tracking nanoparticles at a volume acquisition time of 1 ms using HR-LFM. (a) 3D reconstruction of a representative HGC nanoparticle suspended in water, labeled with Cyanine3. The 3D view was processed by a tricubic-smooth function. (b) Cross-sectional *x*-*y*, *y*-*z*, and *x*-*z* views at *t* = 1 ms across the center (*x*, *y*, *z*) = (0.875 μm, 0.875 μm, 1.000 μm) of the reconstructed volume in (a). The intensity-information is color-coded according to the color scale bar. (c) Time series (*t* = 1 to 10 ms) of the lateral (*x*-*y*) cross-sectional views of the nanoparticle. The depth information is color-coded according to the color scale bar. (d) 3D trajectory of the nanoparticle moving below the diffraction limit (<300 nm). Different time-points (*t* = 1 to 10 ms) are color-coded according to the color scale bar. Scale bars, 500 nm (b, c).

Finally, we tracked HGC nanoparticles suspended in water, labeled with a fluorescent dye Cyanine3 (cy3; **Fig. 6**). HGC nanoparticles were prepared through the covalent attachment of 5β-cholanic acid to glycol chitosan using a previously published protocol [22]. Specifically, 150 mg of 5β-cholanic acid dissolved in 60 ml of methanol was activated with 1.5 mol equivalents of N-hydroxysulfosuccinimide (NHS, ThermoFisher) and 1-ethyl-3-(3- dimethylaminopropyl) carbodiimide hydrochloride (EDC, ThermoFisher). The activated 5β cholanic solution was slowly added to the glycol chitosan solution (500 mg/60 ml, in HPLC water) and stirred for 24 hours at room temperature to ensure complete reaction. The resulting mixture was dialyzed using 10 kDa molecular weight cut-off dialysis cassettes for 24 hours against a water-methanol mixture (1:4 vol/vol), for another 24 hours against water, and lyophilized. For labeling, cy3 dye (Lumiprobe, 1 mg) was dissolved in dimethyl sulfoxide (DMSO, 200 μl) and added dropwise to HGC (100 mg/40 ml, in DMSO) under gentle stirring at room temperature for 6 hours in darkness. The mixture was dialyzed using 3.5 kDa molecular weight cut-off dialysis cassettes for 2 days against HPLC water, and lyophilized. The cy3-HGC nanoparticles were suspended in HPLC water at a concentration of 1 mg/ml and treated with a probe-type sonicator (S-450D Sonifier, Branson Ultrasonics) at 90 W for 6 mins. One drop (5 μl) of cy3-HGC suspension was added to a microscope slide and coverslipped prior to imaging.

At a volume acquisition time of 1 ms, single nanoparticles moving up to 92.81 μm s^−1^ have been recorded using HR-LFM (**Fig. 6a-c** and **Visualization 5**). The 3D positions and trajectories of the nanoparticles were determined by localizing the reconstructed particles using Gaussian fitting with nanometer-level precision in all three dimensions [23] (**Fig. 6d**). The measurements of the nanoparticles are consistent with the reconstructed PSF values using the fluorescent beads (**Fig. 2b**and **Appendix 4**), showing no compromise in spatial precision as the acquisition is accelerated. The system thus has demonstrated the capability of recording dynamic particle behavior in a volumetric context, which has been a spatio-temporal-limiting step for live-cell imaging.

## Conclusion

In summary, we have developed HR-LFM for volumetric live-cell imaging with a spatial resolution of 300-700 nm in all three dimensions, an imaging depth of several micrometers, and a volume acquisition time on the order of milliseconds. Defocusing the MLA effectively mitigates the prohibitive reconstruction artifacts in the previous LFM design, providing four to five-fold larger DOF than conventional high-resolution wide-field microscopy. In addition, due to its greater axial sensitivity and discrimination, we demonstrated the remarkable resolution improvement especially in the axial dimension (i.e. near-isotropic) within a substantial axial range. These findings may lead to new imaging physics and applications for MLA-facilitied microscopy. Advancing current LFM to the subcellular level, the system enables high-speed, volumetric visualization of dynamics and structures in single-cell specimens with low photodamage. Combining the molecular specificity of fluorescent labeling, great scalability, engineered MLAs and deconvolution algorithms, LFM is becoming a particularly promising tool for imaging diverse anatomical and functional traits, spanning molecular, cellular and tissue levels.

## Appendices

## Appendix 1: Details on system alignment

The sCMOS camera was first installed on the camera port of the microscope, and the position of the objective lens was adjusted to form an in-focus image of the sample. Next, the 1:1 relay lens was mounted onto the camera. The camera was placed on a dovetail optical rail (RLA150/M, Thorlabs), aligned to the optical axis, and translated until the same in-focus wide-field image was captured. At this point, the camera was conjugated to the NIP. The MLA, mounted in a translational stage, was first inserted near the NIP where the modulation of the MLA (edges of the microlenses) can be observed on the camera. The MLA was then slightly adjusted until the modulation of the MLA disappeared on the camera. At this point, the MLA was aligned to the NIP, and the camera recorded the wide-field images reported in this work. For light-field imaging, the camera was translated away from the sample by the focal length of the MLA (*f*_*ml*_ = 3.75 mm), where the system formed conventional LFM (**Appendix 2**; *a* = 0 and *b* = *f*_*ml*_ = 3.75 mm). To establish the design in this work, the MLA was translated away from the sample by *a* = 25 mm, and the camera, accordingly, by *a* + *b* - *f*_*ml*_ = 25.25 mm. At this point, the HR-LFM system was established as reported (*a* = 25 mm and *b* = 4 mm) (**Fig. 1a** and **Appendix 2**).

## Appendix 2: Design schematics of conventional LFM, focused plenoptic camera, and HR-LFM

**Fig. 7.**
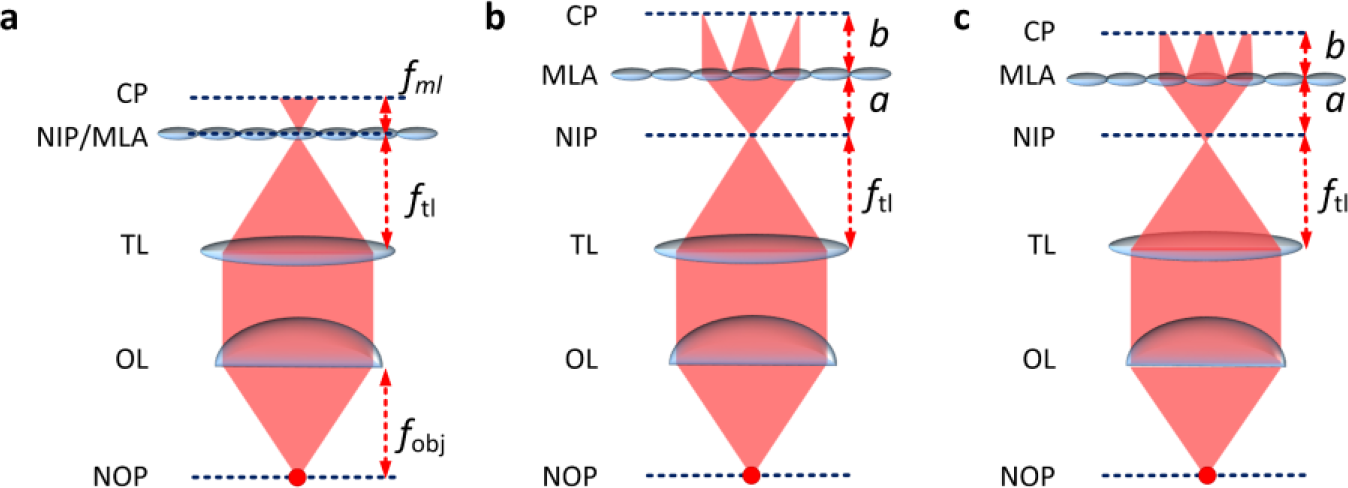
Design schematics of conventional LFM (a), focused plenoptic camera (b), and HR-LFM (c). (a) For conventional LFM, the MLA is placed at the NIP of a wide-field microscope (*a* = 0 and *b* = *f*_*ml*_) [1,2]. Near the NIP, the sampling pattern of the spatial information becomes redundant (i.e. restrained to one microlens). As a result, the wave-optics based model produces prohibitive reconstruction artifacts across the NIP. (b) For the focused plenoptic camera, the MLA forms an imaging relationship between the NIP and the camera (1/*a* + 1/*b* = 1/*f*_*ml*_) [17]. The sampling geometry of the angular information becomes less widely distributed, inherently impairing the refocusing (or volumetric imaging) capability. (c) For HR-LFM, the MLA is placed at a defocused position (1/*a* + 1/*b* > 1/*f*_*ml*_) to facilitate simultaneous, dense sampling of both spatial and angular information. *f*_*ml*_, the focal length of the MLA; *f*_tl_, the focal length of the tube lens; *f*_obj_, the effective focal length of the objective lens; *a* and *b* denote the distances from the MLA to the NIP and the camera, respectively; TL, tube lens; OL, objective lens; NOP, native object plane; CP, camera plane.

## Appendix 3: Model of light-field propagation and image formation

Projecting the 3D volume in the object domain to the 2D imaging space, the wavefunction at the NIP using the high-NA objective lens, is predicted by the Debye theory as [18]:

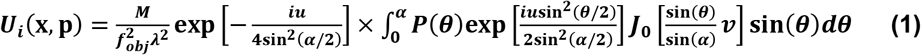

where ***f***_*obj*_ is the focal length of the objective lens, and ***J***_0_ is the zeroth order Bessel function of the first kind. The variables **𝒗** and ***u*** represent normalized radial and axial coordinates; the two variables are defined by **𝒗** = ***k***[(***x***_1_ − ***p***_1_)^2^ + (***x***_2_ − ***p***_2_)^2^]^1/2^ ***sin***(***α***)and ***u*** = 4*kp*_3_sin^2^ (*α*/2); **p** = (***p*_1_, *p*_2_, *p*_3_**) is the position for a point source in a volume in the object domain; **x** = (***x*_1_, *x*_2_**) ∈ ***R*^2^** represents the coordinates on the NIP; ***M*** is the magnification of the objective lens; the half-angle of the NA is ***α* = sin^−1^(*NA*/*n***); the wavenumber ***k* =2*πn*/*λ*** were calculated using the wavelength ***λ*** and the refractive index ***n*** of the immersion medium. For Abbe-sine corrected objective lenses, the apodization function of the microscope ***p*(*θ*)=cos(*θ*)^1/2^** was used.

When the MLA is located by the distance ***a*** from the NIP, the wavefront on the MLA can be represented by Fresnel propagation from the NIP, given as:

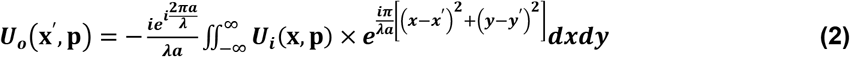

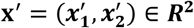 represents the coordinates on the MLA. The aperture of a microlens can be described as an amplitude mask **rect**(**x**′/***d***);, combined with a phase mask 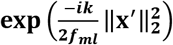.

The modulation induced by a microlens is then described as:

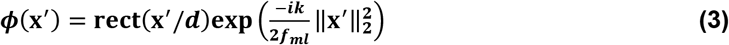

where ***f_ml_*** is the focal length of the MLA, and ***d*** is the pitch of the MLA (or the diameter of a single microlens). Thus, the modulation of the entire MLA, composed of periodic microlenses, can be described by convolving ***ϕ***(**x**′) with a 2D comb function **comb**(**x**′/***d***), i.e. ***Φ***(**x**′) = **ϕ**(**x**′)⨂**comb**(**x**′/***d***).

Next, the light propagation over the distance of ***b*** from the MLA to the camera can be modelled using Fresnel propagation. The final complex-valued PSF is described as:

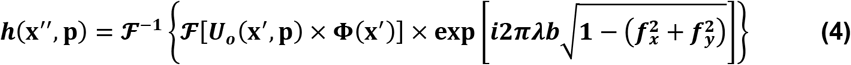

where the exponential term is the Fresnel transfer function, ***f*_*x*_** and ***f*_*y*_** are the spatial frequencies in the camera plane, and 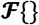 and 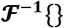 represent the Fourier transform and inverse Fourier transform operators, respectively. The final intensity image ***o*(x″)** at the camera plane is described by

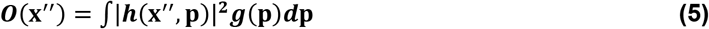

where, as previously defined, **p ∈ *R*^3^** is the position in a volume containing isotropic emitters, whose combined intensities are distributed according to ***g*(p)**. In the discrete model of the complex-valued PSF, ***h*(x″, p)** is represented by the measurement matrix **H** which elements *h*_*kj*_ represent the projection of the light arriving at the pixel ***o*(*j*)** on the camera from the *k*^th^ voxel ***g*(*k*)** in the object space [3]. The volume reconstruction utilized a deconvolution algorithm based on the inverse problem in tomographic image formation [19], where the volumetric information was obtained from multiple different perspectives of a 3D volume using deconvolution. The algorithm was further modified combining the wave-optics model for reconstruction of light-field data [3,4] (**Supplementary Code**).

## Appendix 4: Full-width at half-maximum (FWHM) values of the reconstructed point spread functions (PSFs) at varying depths

**Fig. 8.**
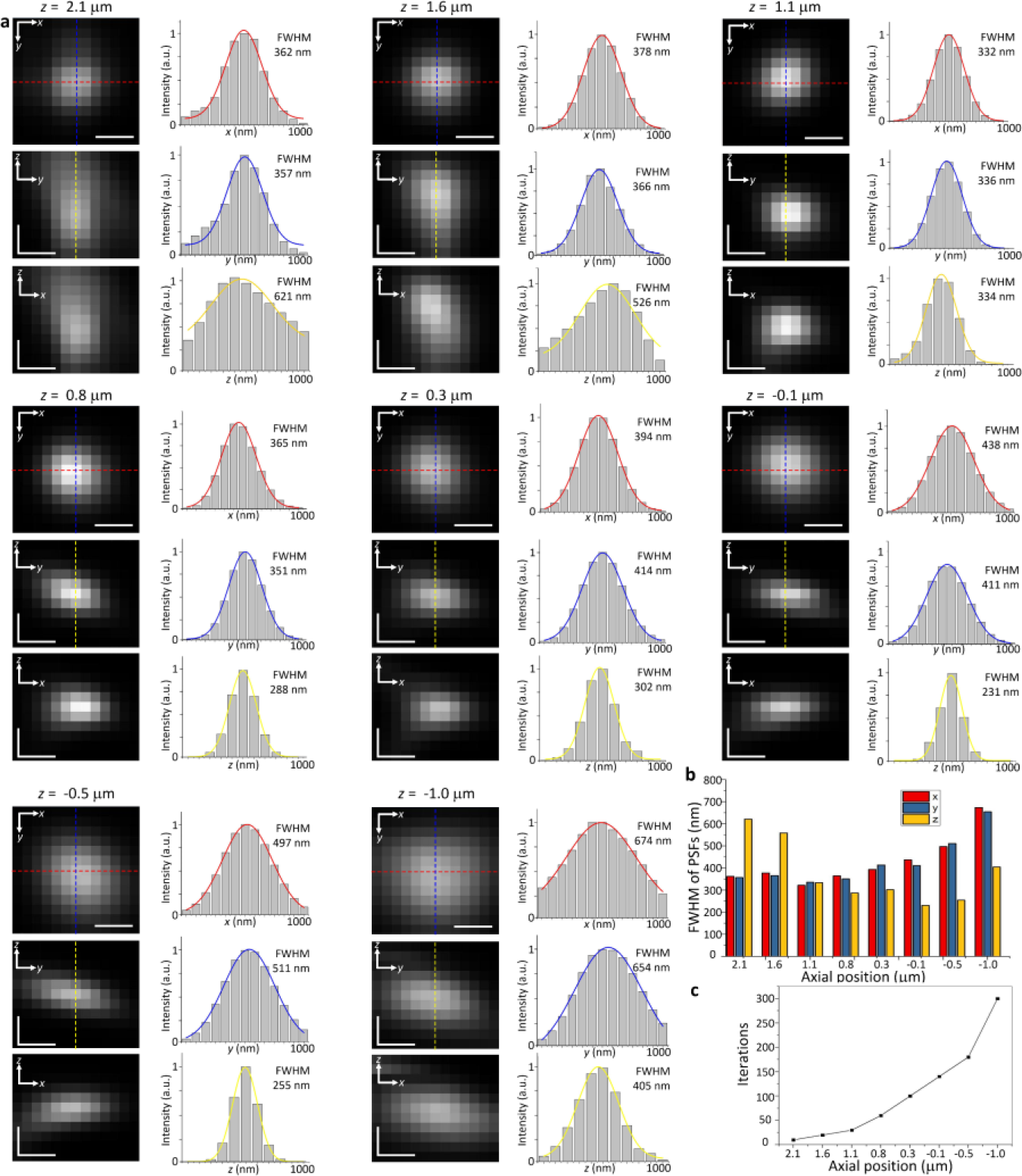
Full-width at half-maximum (FWHM) values of the reconstructed point-spread functions (PSFs) at varying depths. (a) The reconstructed cross-sectional images (left columns) and their corresponding profiles along the dashed lines (right columns) of sub-diffraction-limited 100-nm fluorescent beads in lateral (*x*-*y*) and axial (*x*-*z* and *y*-*z*) dimensions at depths (*z* = 2.1, 1.6, 1.1, 0.8, 0.3, −0.1, −0.5, and −1.0 μm), respectively. (b) The 3D FWHM values of the profiles at varying depths using Gaussian fitting. The reconstructed PSFs exhibited a near-diffraction limited 300-700 nm resolution in all three dimensions over a >3 μm range. Detailed descriptions of the axial-resolving capability are demonstrated in **Appendix 5**. (c) Numbers of iterations taken for the reconstruction at varying depths. The numbers were determined based on the distribution of the optical signals across the MLA and the corresponding signal-to-noise ratio (SNR). These numbers were consistently used in this work for other samples. Scale bars, 300 nm.

## Appendix 5: Improved axial-resolving capability of HR-LFM

**Fig. 9.**
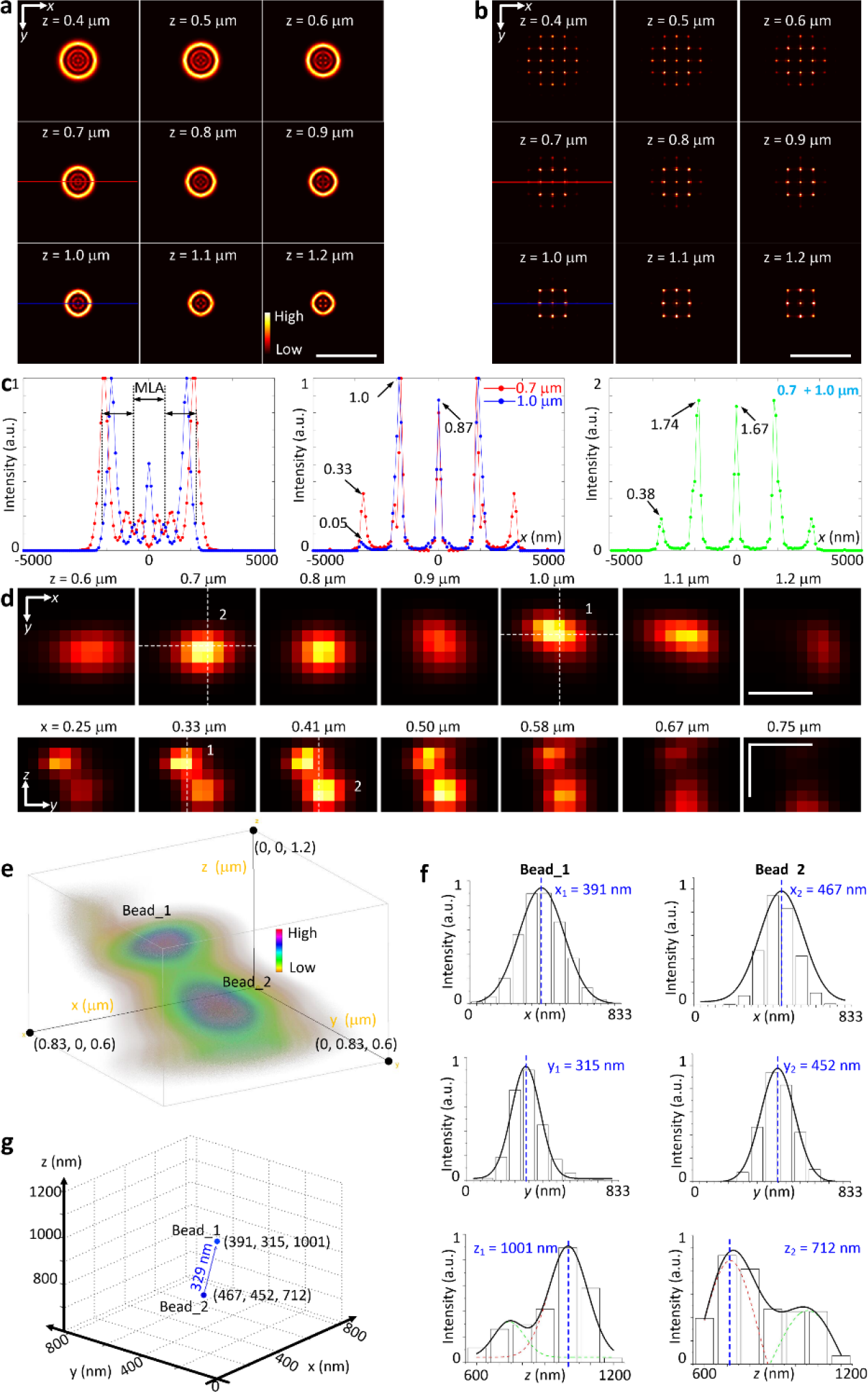
Improved axial-resolving capability of HR-LFM. (a,b) Patterns of the optical signals entering the MLA (a) and at the camera sensor (b) for PSFs at varying depths. Notably, the MLA can segmentally reveal the angular information in the wavefront (i.e. spatial frequencies) of the optical signals, which provides higher sensitivity against fine axial variations. As seen in (b), the PSF of HR-LFM changes significantly for an axial displacement of 300 nm (e.g. at *z* = 0.7 μm and 1.0 μm), which distribution varies from covering primarily 5 × 5 to 3 × 3 microlenses, respectively. However, such variation was less remarkable in the standard Gaussian PSF (a) over the same displacement of 300 nm. (c) The cross-sectional profiles of the standard Gaussian (left) and the HR-LFM (middle) PSFs at *z* = 0.7 μm and 1.0 μm, respectively. As seen, the standard Gaussian PSF was moderately expanded by a distance (~250 nm), which is below the diameter of a single microlens (effectively 125 μm / 100 = 1.25 μm in the object domain). In contrast, such an expansion was better recognized by HR-LFM, resulting in a significantly changed intensity pattern on the camera (e.g. at *z* = 0.7 μm and 1.0 μm, the number of major intensity peaks reduces from 5 to 3, respectively). In addition, at each depth, the corresponding PSF of HR-LFM exhibited distinct ratios between its major intensity peaks. When there are multiple PSFs situated at different axial positions, the camera captures the sum of these PSFs. For example, the *right* figure in (c) shows the summed intensity pattern of two PSFs located at the same lateral position but different axial positions at *z* = 0.7 μm and 1.0 μm. While overlapping in the raw data, individual PSFs can be decoupled with the wave-optics based model which convergence considers the prior knowledge of individual PSF patterns at varying depths through iterative deconvolution. (d) Experimental results for imaging two nearby 100-nm fluorescent beads using HR-LFM. The top panel shows the reconstructed lateral cross-sectional images of two resolved beads (1 and 2) at varying depths (left to right, *z* = 0.6, 0.7, 0.8, 0.9, 1.0, 1.1, and 1.2 μm, respectively). The bottom panel shows the reconstructed axial cross-sectional images in *y*-*z* of the two resolved beads at varying *x*-positions (left to right, *x* = 0.25, 0.33, 0.41, 0.50, 0.58, 0.67, and 0.75 μm, respectively). (e) 3D rendering of the reconstructed beads. (f,g) Cross sectional profiles and positions of the two beads in three dimensions. The 3D positions of the beads and the FWHM values of their profiles were determined using Gaussian fitting. As measured, the distance between the two beads was 329 nm, slightly below the diffraction limit in the axial dimension (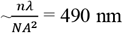, where the refractive index of immersion oil n = 1.515, the emission wavelength λ = 680 nm, and NA = 1.45). This resolving capability was achieved due to the sensitive angular detection using the MLA in HR-LFM. Scale bars, 5 μm (a,b), 500 nm (d).

## Appendix 6: Imaging mitochondria in Drp1^−/−^ mouse embryo fibroblasts (MEFs) using HR-LFM

**Fig. 10.**
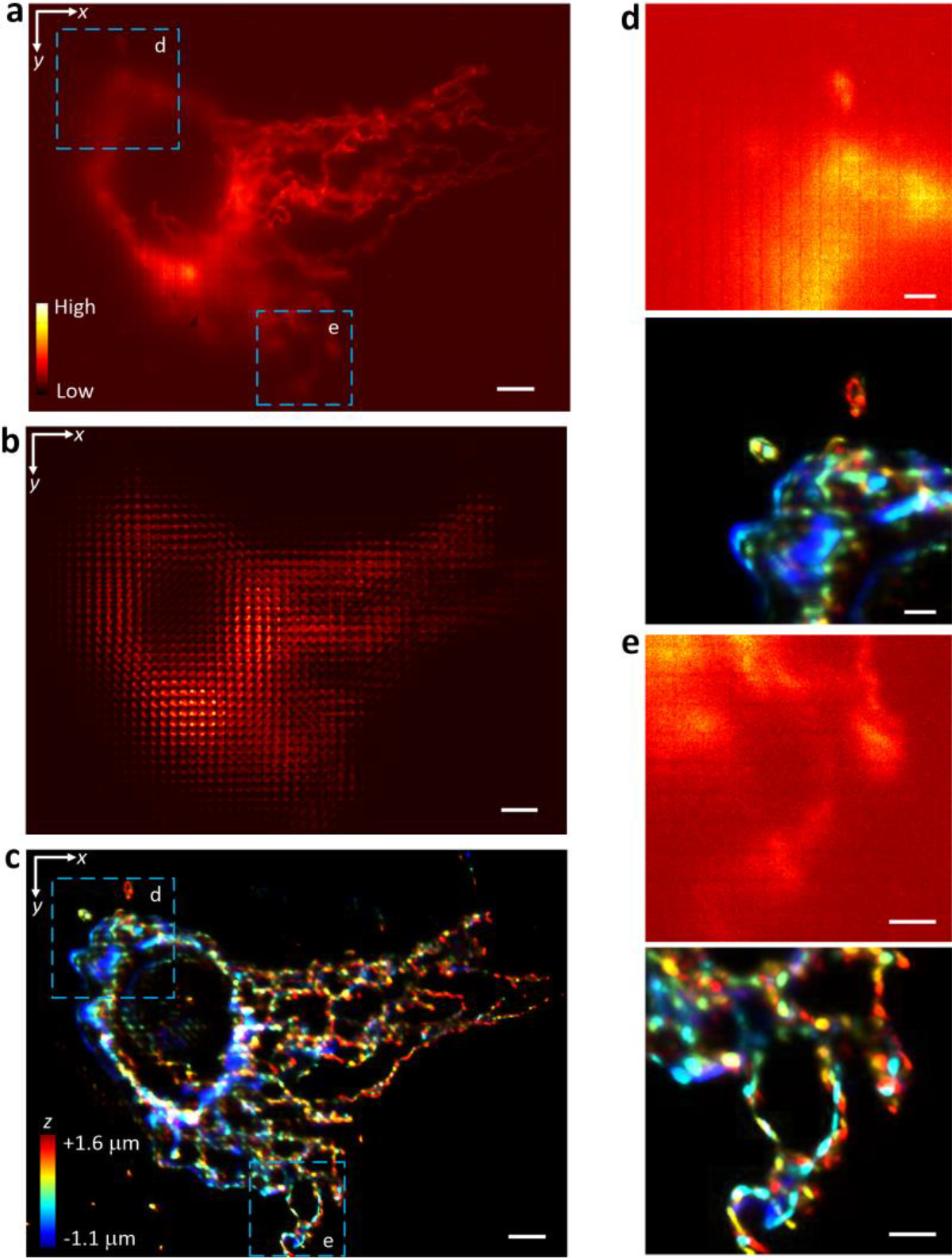
Imaging mitochondria in *Drp1* −/− mouse embryo fibroblasts (MEFs) using HR-LFM. (a-c) Conventional wide-field image (a), light-field raw image (b), and reconstructed 3D image (c) of mitochondria in *Drp1*^−/−^ MEFs labeled with MitoTracker acquired with a volume acquisition time of 0.1s and at an axial step size of 100 nm. The depth-information is color coded according to the color scale bar in (c). (d,e) Zoomed-in images of the boxed regions in (a) (top) and the corresponding boxed regions in (c) (bottom), respectively, showing the high spatial resolution and volumetric imaging capability of HR-LFM. Scale bars, 5 μm (a-c), 2 μm (d,e).

## Appendix 7: Imaging Golgi complex in HeLa cells using HR-LFM

**Fig. 11.**
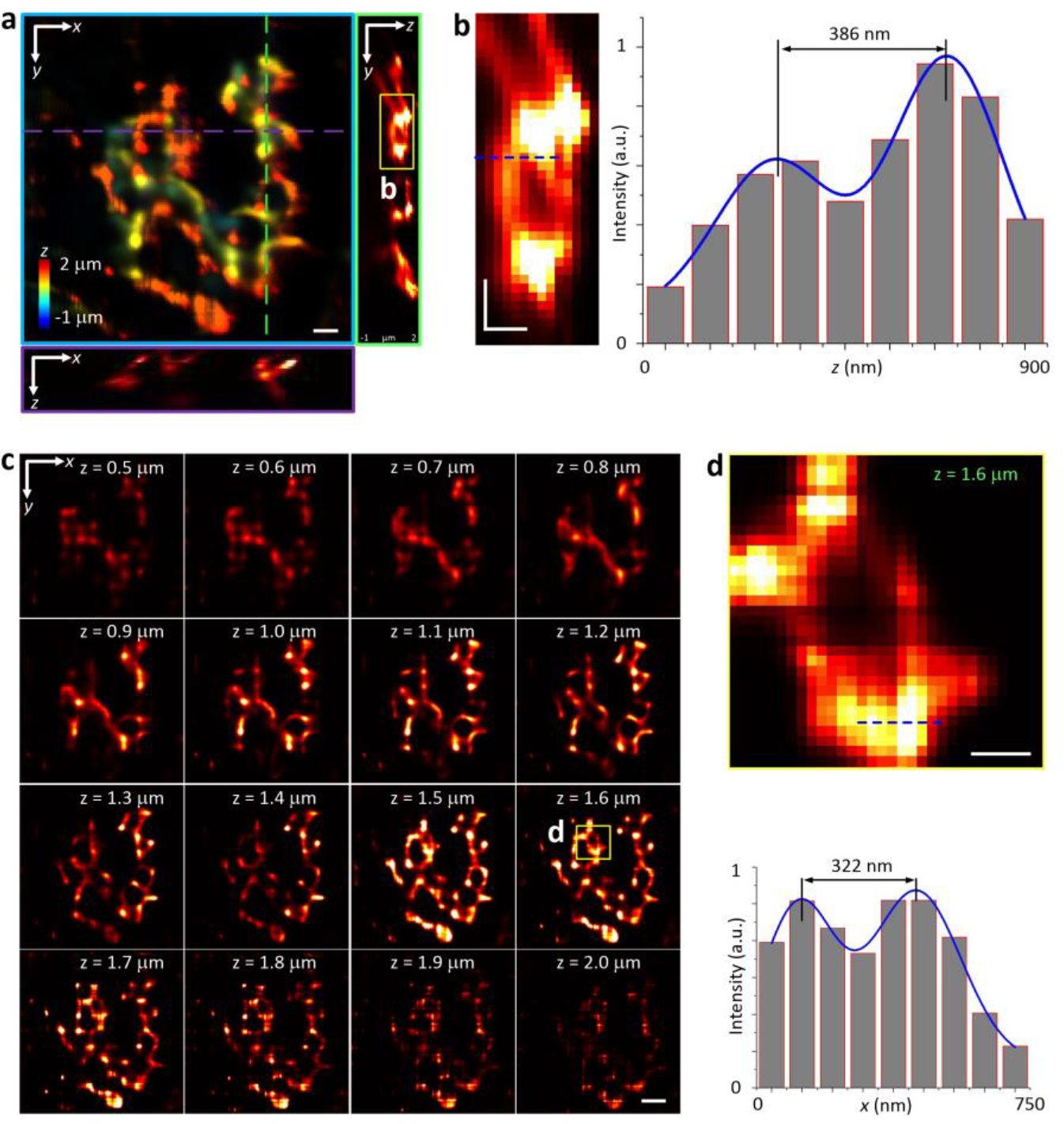
Imaging Golgi complex in HeLa cells using HR-LFM. (a) 3D reconstruction of the Golgi complex in HeLa cells. The Golgi structures were visualized using immunofluorescence staining for the Golgi marker GM130 with DyLight 549. The depth-information is color-coded according to the color scale bar. The insets show the cross-sectional images along the dashed lines in *x*-*z* and *y*-*z*. (b) *Left*, the zoomed-in image of the boxed region in (a), showing axially resolved inner structures in the Golgi complex. *Right*, the cross-sectional profile along the dashed line, showing that axial structures as close as ~400 nm can be resolved. (c) Reconstructed *z*-stack images at an axial step size of 100 nm, showing fine structural variations captured at various axial positions. (d) *Top*, zoomed-in image of the boxed region in (c) at *z* = 1.6 μm. *Bottom*, the cross-sectional profile along the dashed line shows that lateral structures as close as ~400 nm can be resolved. The measurements are consistent with the FWHM values in **Appendix 4**. Scale bars, 1 μm (a), 500 nm (b,d), 2 μm (c).

## Appendix 8: Imaging Golgi-derived membrane vesicles in living COS-7 cells using HR-LFM

**Fig. 12.**
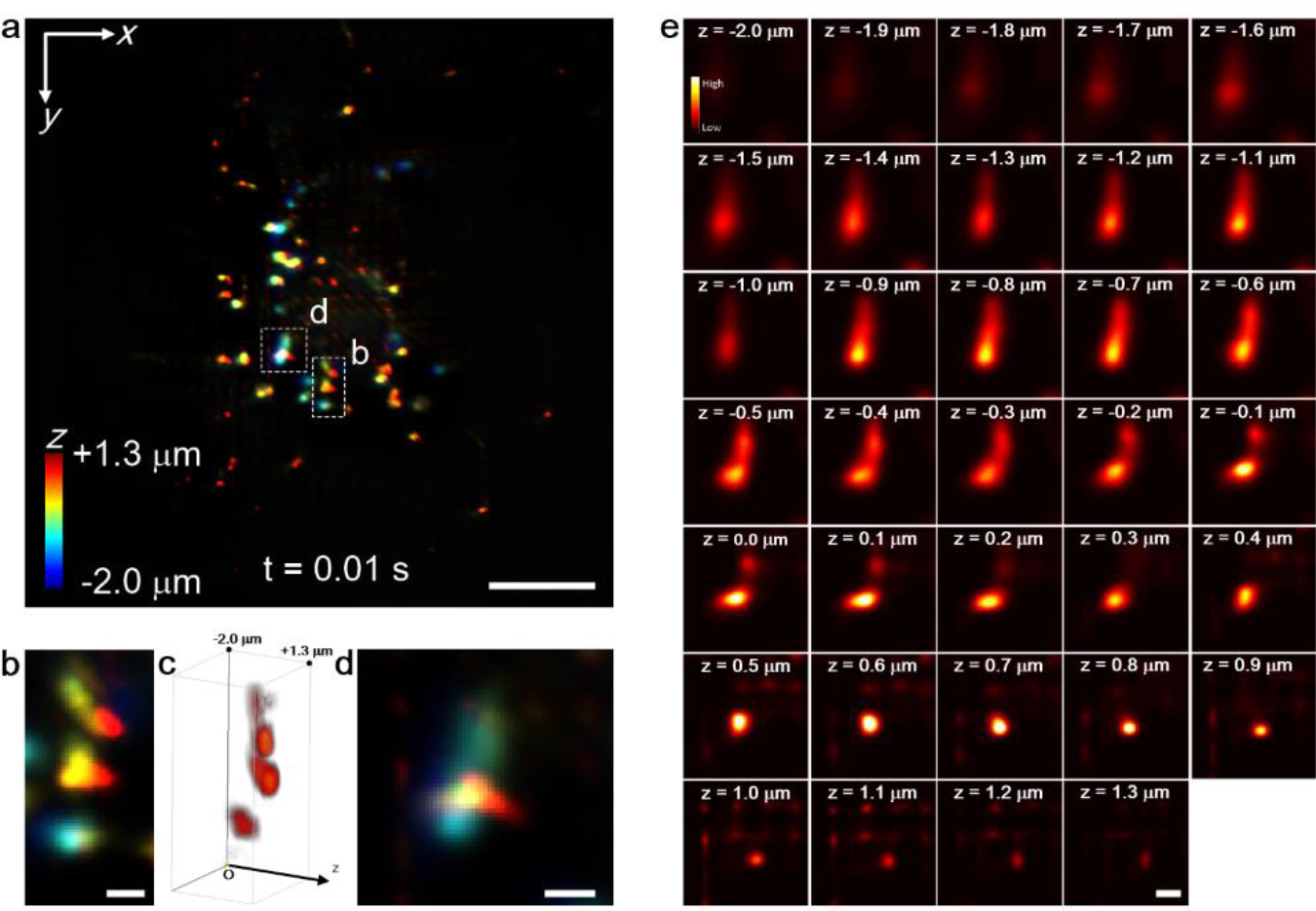
Imaging Golgi-derived membrane vesicles in living COS-7 cells using HR-LFM. (a) Reconstructed 3D image of vesicles in living COS-7 cells labeled with mEmerald-Golgi-7 acquired with a volume acquisition time of 0.01 s. The depth-information (over a >3 μm range) is color-coded according to the color scale bar. (b) Zoomed-in image of the corresponding boxed region in (a). (c) 3D view of (b) processed by a tricubic-smooth function. (d) Zoomed-in image of the corresponding boxed region in (a). (e) *z*-stack images of (d) at an axial step size of 100 nm from *z* = −2.0 μm to +1.3 μm. (b-e) show that several vesicles separated less than 1 μm were resolved in all three dimensions. Scale bars: 10 μm (a), 1 μm (b,d,e).

## Appendix 9: Imaging Golgi-derived membrane vesicles in living COS-7 cells at a volume acquisition time of 5 ms using HR-LFM

**Fig. 13.**
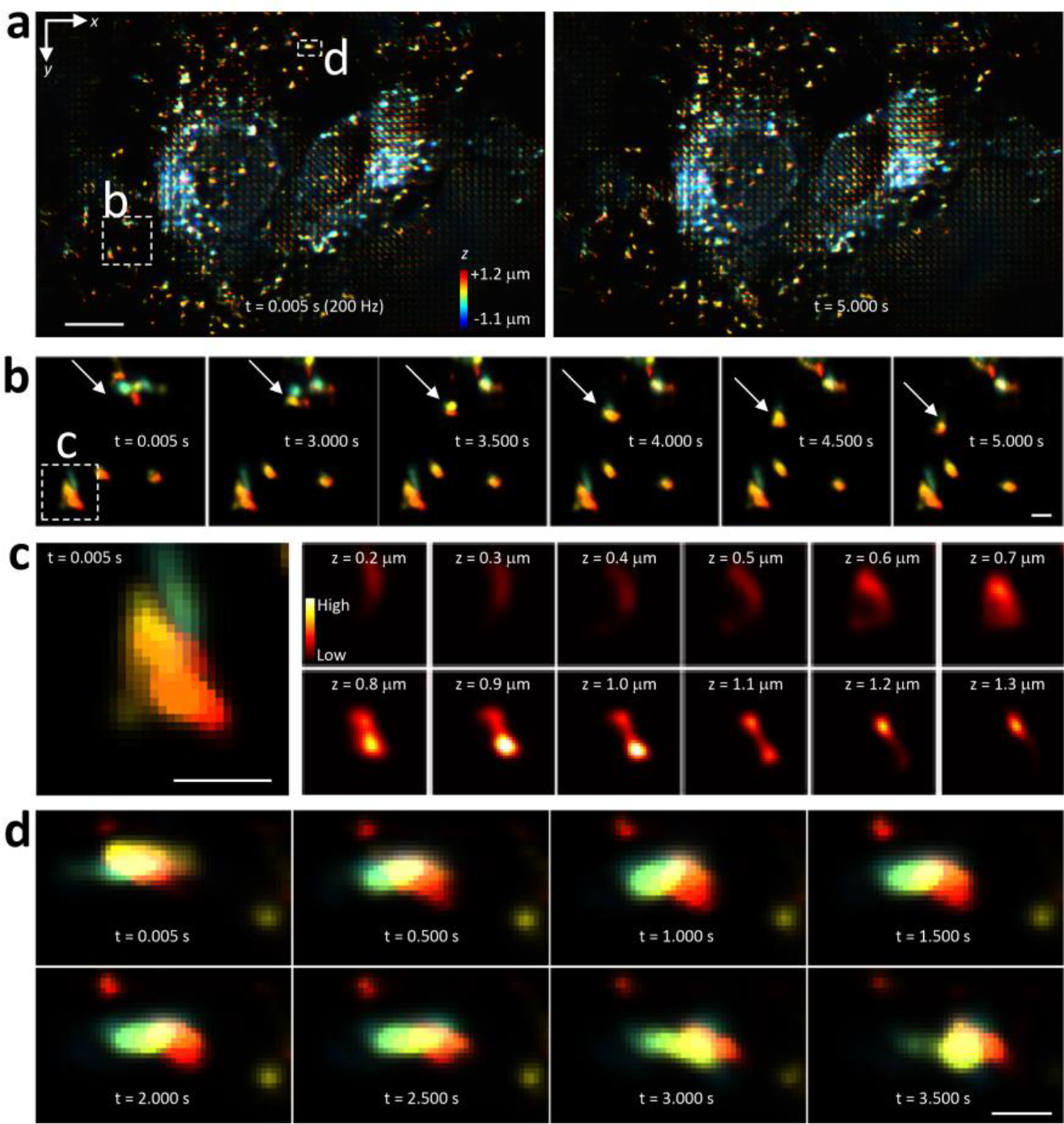
Imaging Golgi-derived membrane vesicles in living COS-7 cells at a volume acquisition time of 5 ms using HR-LFM. (a) Reconstructed 3D images of vesicles in living COS-7 cells labeled with mEmerald-Golgi-7 acquired at a volume acquisition time of 5 ms. The left and right are reconstructed 3D images at *t* = 0.005s and 5.000s of a 1000-time-point series, respectively. The depth-information is color-coded according to the color scale bar. (b) Zoomed in images of the corresponding boxed region in (a) at *t* = 0.005s, 3.000s, 3.500s, 4.000s, 4.500s and 5.000s, respectively. White arrows indicate vesicles moving during time-points. (c) Zoomed-in image (leftmost) of the boxed region in (b) at *t* = 0.005s and its *z*-stack images from *z* = 0.2 μm to +1.3 μm at an axial step size of 100 nm, resolving several nearby vesicles. (d) Zoomed-in images of the corresponding boxed region in (a) at *t* = 0.005s, 0.500s, 1.000s, 1.500s, 2.000s, 2.500s, 3.000s and 3.500s, respectively. Scale bars: 10 μm (a), 1 μm (b-d).

## Appendix 10: Acquisition parameters for all images in Figs. 1-6, 8-13

| Image Property | Fig. 1e | Fig. 2a | Fig. 2c, e | Fig. 3 |
| --- | --- | --- | --- | --- |
| sample | 100-nm bead | 100-nm bead | 1-μm microsphere | HeLa (fixed) |
| supplier | T7279, ThermoFisher | T7279, ThermoFisher | F14791, ThermoFisher | ATCC |
| fluorescent label | Blue, Dark Red, Green, Orange | Blue, Dark Red, Green, Orange | Dark Red, Green, Orange | AlexaFluor 647 / anti-Tom20 (mito) |
| excitation wavelength | 647 nm | 647 nm | 647 nm | 647 nm |
| emission wavelength | 680 nm | 680 nm | 680 nm | 667 nm |
| dichroic mirror | T660lpxr, Chroma | T660lpxr, Chroma | T660lpxr, Chroma | T660lpxr, Chroma |
| emission filter | ET700/75, Chroma | ET700/75, Chroma | ET700/75, Chroma | ET700/75, Chroma |
| FOV (μm<sup>2</sup>) | 133 x 133 | 133 x 133 | 133 x 133 | 133 x 133 |
| sCMOS frame rate / volume acquisition time | 100 ms | 100 ms | 100 ms | 100 ms |
| Image Property | Fig. 4 | Fig. 5 | Fig. 6 |  |
| sample | Drp1<sup>-/-</sup> Mouse embryo fibroblasts (live) | COS-7 (live) | Hydrophobically-modified glycol chitosan (HGC) nanoparticles |  |
| supplier | Hiromi Sesaki (JHU) | ATCC | Yizhi Meng (SBU) |  |
| fluorescent label | MitoTracker Deep Red | mEmerald-Golgi-7 | Cyanine3 |  |
| excitation wavelength | 647 nm | 488 nm | 561 nm |  |
| emission wavelength | 680 nm | 518 nm | 570 nm |  |
| dichroic mirror | T660lpxr, Chroma | T495lpxr, Chroma | T560lpxr, Chroma |  |
| emission filter | ET700/75, Chroma | ET525/50, Chroma | FF02-617/73, Semrock |  |
| FOV (μm<sup>2</sup>) | 133 x 133 | 133 x 133 | 8.3 x 133 | |
| sCMOS frame rate / volume acquisition time | 10 Hz / 100 ms | 100 Hz / 10 ms | 1000 Hz / 1 ms |  |
| Image Property | Fig. 8 | Fig. 9 | Fig. 10 | Fig. 11 |
| sample | 100-nm bead | 100-nm bead | Drp1<sup>-/-</sup> Mouse embryo fibroblasts (live) | HeLa (fixed) |
| supplier | T7279, ThermoFisher | T7279, ThermoFisher | Hiromi Sesaki (JHU) | ATCC |
| fluorescent label | Blue, Dark Red, Green, Orange | Blue, Dark Red, Green, Orange | MitoTracker Deep Red | DyLight 549 / GM130 (Golgi) |
| excitation wavelength | 647 nm | 647 nm | 647 nm | 561 nm |
| emission wavelength | 680 nm | 680 nm | 680 nm | 576 nm |
| dichroic mirror | T660lpxr, Chroma | T660lpxr, Chroma | T660lpxr, Chroma | T560lpxr, Chroma |
| emission filter | ET700/75, Chroma | ET700/75, Chroma | ET700/75, Chroma | FF02-617/73, Semrock |
| FOV (μm<sup>2</sup>) | 133 x 133 | 133 x 133 | 133 x 133 | 133 x 133 |
| sCMOS frame rate / volume acquisition time | 10 Hz / 100 ms | 10 Hz / 100 ms | 10 Hz / 100 ms | 10 Hz / 100 ms |
| Image Property | Fig. 12 | Fig. 13 |  |  |
| sample | COS-7 (live) | COS-7 (live) |  |  |
| supplier | ATCC | ATCC |  |  |
| fluorescent label | mEmerald-Golgi-7 | mEmerald-Golgi-7 |  |  |
| excitation wavelength | 488 nm | 488 nm |  |  |
| emission wavelength | 518 nm | 518 nm |  |  |
| dichroic mirror | T495lpxr, Chroma | T495lpxr, Chroma |  |  |
| emission filter | ET525/50, Chroma | ET525/50, Chroma |  |  |
| FOV (μm<sup>2</sup>) | 133 x 133 | 67 x 133 | | |
| sCMOS frame rate / volume acquisition time | 100 Hz / 10 ms | 200 Hz / 5 ms |  |  |

## Appendix 11: Additional parameters for all Visualizations 1-5

|  |  |
| --- | --- |
| **Visualization 1** | Time-lapse video of reconstructed mitochondria in living *Drp1*<sup>-/-</sup> MEFs (FOV = 133 x 133 μm, volume acquisition rate = 10 Hz). Scale bars, 10 μm (main), 1 μm (insets). Color scale bar, z = -1.0 μm (blue) to 1.5 μm (red). |
| **Visualization 2** | Zoomed-in, 3D view of the corresponding region (marked with 'triangle') in Supplementary Video 1 (volume acquisition rate = 10 Hz). |
| **Visualization 3** | Zoomed-in, 3D view of the corresponding region (marked with 'cross') in Supplementary Video 1 (volume acquisition rate = 10 Hz). |
| **Visualization 4** | Time-lapse video of reconstructed Golgi-derived membrane vesicles in living COS-7 cells (FOV = 133 x 133 μm, volume acquisition rate = 100 Hz). Scale bars, 10 μm (main), 1 μm (insets). Color scale bar, z = -1.5 μm (blue) to 1.0 μm (red). |
| **Visualization 5** | Time-lapse video of a reconstructed diffusing nanoparticle suspended in water (FOV = 8.3 x 133 μm, volume acquisition rate = 1000 Hz). |

## Funding

National Institutes of Health grants 1R35GM124846 (to S.J.) and R01GM084251 (to M.F.), National Science Foundation grants CBET1604565 and EFMA1830941 (to S.J.).

## Acknowledgments

We acknowledge the support of the NSF-CBET Biophotonics program, the NSF-EFMA program, and the NIH-NIGMS MIRA program.

## Disclosures

The authors declare that there are no conflicts of interest related to this article.

